# The Immune-Modulatory Function of Megakaryocytes in the Hematopoietic Niche of Myeloproliferative Neoplasms

**DOI:** 10.1101/2025.04.01.646152

**Authors:** Sandy Lee, Xiaoxi Yang, Kyla Masarik, Tameena Ahmed, Lei Zheng, Huichun Zhan

## Abstract

Myeloproliferative neoplasms (MPNs) are clonal stem cell disorders characterized by dysregulated megakaryopoiesis and neoplastic hematopoietic stem cell (HSC) expansion. Using a murine model with MK-specific JAK2V617F expression, we establish an MPN aging model where mutant MKs drive HSC expansion and a progressive decline in wild-type HSC function. Compared to wild-type MKs, JAK2V617F MKs exhibit heightened inflammation and innate immune activation with aging, including increased antigen presentation, elevated pro-inflammatory cytokines, skewed T cell populations, and impaired T cell functions in the marrow niche. Enhanced MK immunomodulatory function is linked to mutant cell expansion and MPN progression in a chimeric murine model with co-existing wild-type and JAK2V617F mutant HSCs. LINE-1 (long-interspersed element-1), a retrotransposon linked to innate immune activation and aging, is upregulated in mutant MKs during aging in murine models. We validated that LINE-1–encoded protein ORF1p is expressed in marrow MKs in 12 of 13 MPN patients but absent in control samples from patients undergoing orthopedic surgery (n=5). These findings suggest that MKs reprogram the marrow immune microenvironment, impairing normal HSC function while promoting neoplastic expansion in MPNs. LINE-1 activation in mutant MKs may be a key driver of immune dysregulation in MPNs.

**Key Points:**

- JAK2V617F mutant MKs reprogram the marrow immune microenvironment to promote neoplastic HSC expansion in MPNs.
- LINE-1 activation in diseased MKs triggers chronic inflammation and immune dysfunction in MPNs.

## Introduction

Megakaryocytes (MKs), traditionally recognized as the precursors of platelets, have emerged as key regulators of the hematopoietic stem cell (HSC) function. Through the production of cytokines, chemokines, and extracellular matrix components, MKs influence both steady-state and stress hematopoiesis^1-6^. Recent studies also suggest that MKs contribute to pathogen surveillance and immune regulation^7-12^. While abnormal megakaryopoiesis is a common feature of many hematologic malignancies^13-18^, how diseased MKs alter their immunomodulatory function and impact HSC behavior in normal and neoplastic hematopoiesis remains poorly understood.

Myeloproliferative neoplasms (MPNs), including polycythemia vera (PV), essential thrombocythemia (ET), and primary myelofibrosis (PMF), are clonal stem cell disorders characterized by hematopoietic stem/progenitor cell (HSPC) expansion and an increased risk of transformation to acute leukemia. The acquired kinase mutation JAK2V617F plays a central role in these diseases^19,20^. MK hyperplasia is a hallmark feature of MPNs, with advanced disease often associated with morphologically and functionally altered MKs, raising the possibility that diseased MKs contribute to disease progression^13,16,17,21^. Both MPN incidence and leukemia transformation risk increase significantly with aging^19,20^, suggesting a strong interplay between JAK2V617F-driven hematopoiesis and age-related changes in the marrow microenvironment.

Previously, we established an MPN aging model in which JAK2V617F mutant MKs drive a myeloproliferative syndrome characterized by modest thrombocytosis, increased marrow MKs and HSCs, and a gradual decline in wild-type HSC function during a 2-year follow up.^22,23^. In this study, we investigate the immunomodulatory functions of JAK2V617F mutant MKs during aging and MPN disease progression. We demonstrate that mutant MKs exhibit heightened inflammation and innate immune activation with aging, and that enhanced MK immunomodulatory function is associated with MPN disease progression. Notably, LINE-1 (long-interspersed element-1), a retrotransposon linked to innate immune activation, inflammation, and aging, is upregulated in JAK2V617F MKs in aged mice and detected in marrow MKs in 12 of 13 MPN patients, but absent in age-matched controls (n=5). These findings suggest that JAK2V617F mutant MKs actively shape the marrow immune microenvironment, potentially driving disease progression through chronic inflammation and immune dysfunction.

## Methods

### Experimental mice

All mouse experiments were performed according to protocols approved by the Institutional Animal Care and Use Committee at Stony Brook University. JAK2V617F Flip-Flop (FF1) mice (which carry a Cre-inducible human JAK2V617F gene driven by the human JAK2 promoter) were provided by Radek Skoda (University Hospital Basal, Switzerland). Pf4-Cre mice (which express Cre under the promoter of platelet factor 4; JAX stock #008535) were crossed with the FF1 mice to generate a transgenic mouse line with human JAK2V617F expression in the MK lineage (Pf4^+^FF1^+^). and Tie2-cre mice were obtained from Jackson Laboratory. All mice used were housed in a pathogen-free mouse facility at Stony Brook University. No randomization or blinding was used to allocate experimental groups.

### Patient samples

Archived bone marrow biopsies from MPN patients and age-matched orthopedic surgery controls were obtained from Stony Brook University hospital. Sample collection and analysis were conducted under an Institutional Review Board-approved protocol. A detailed list of patient samples used in this study is provided in Figure 7.

Additional details are provided in the Supplementary Information.

## Results

### Inflammatory and immune activation signatures in JAK2V617F mutant MKs linked to skewed marrow T cell populations during aging

To investigate the impact of a JAK2V617F-bearing MK niche on MPN neoplastic hematopoiesis *in vivo*, we crossed mice with a Cre-inducible human JAK2V617F transgene (FF1)^24^ with the Pf4-cre mice, in which Cre recombinase is driven by the MK-specific platelet factor 4 promoter^25^. Although the Pf4-cre model has some limitations including its expression in other cell lineages, it has remained a gold standard for creating efficient MK-specific transgene expression^26^. Using rigorous and sensitive assays (e.g., expression of fluorescent reporter genes, expression of the JAK2V617F transgene, functional analysis of JAK/STAT downstream signaling), we and others have verified the specific transgene expression and/or activation in MKs, but not in HSCs^3,22,23,27,28^. The resulting Pf4-cre^+^FF1^+^ (Pf4^+^FF1^+^) mice develop an essential thrombocythemia phenotype characterized by modest neutrophilia and thrombocytosis during 2 years of follow up (Figure 1A). Flow cytometry analysis of marrow revealed about 2-fold expansion of CD41^+^ MKs and Lineage^-^cKit^+^Sca1^+^CD150^+^CD48^-^ HSCs in Pf4^+^FF1^+^ mice compared to age-matched control mice^23^. In addition, we found that the 2-year-old Pf4^+^FF1^+^ mice exhibit many features of HSC impairment similar to aging, including reduced engraftment capacity, myeloid-skewed hematopoiesis with expanded CD41^+^ myeloid-biased HSCs, and decreased HSC quiescence^23^. These observations suggest that JAK2V617F mutant MKs progressively impair HSC function with aging.

**Figure 1.**
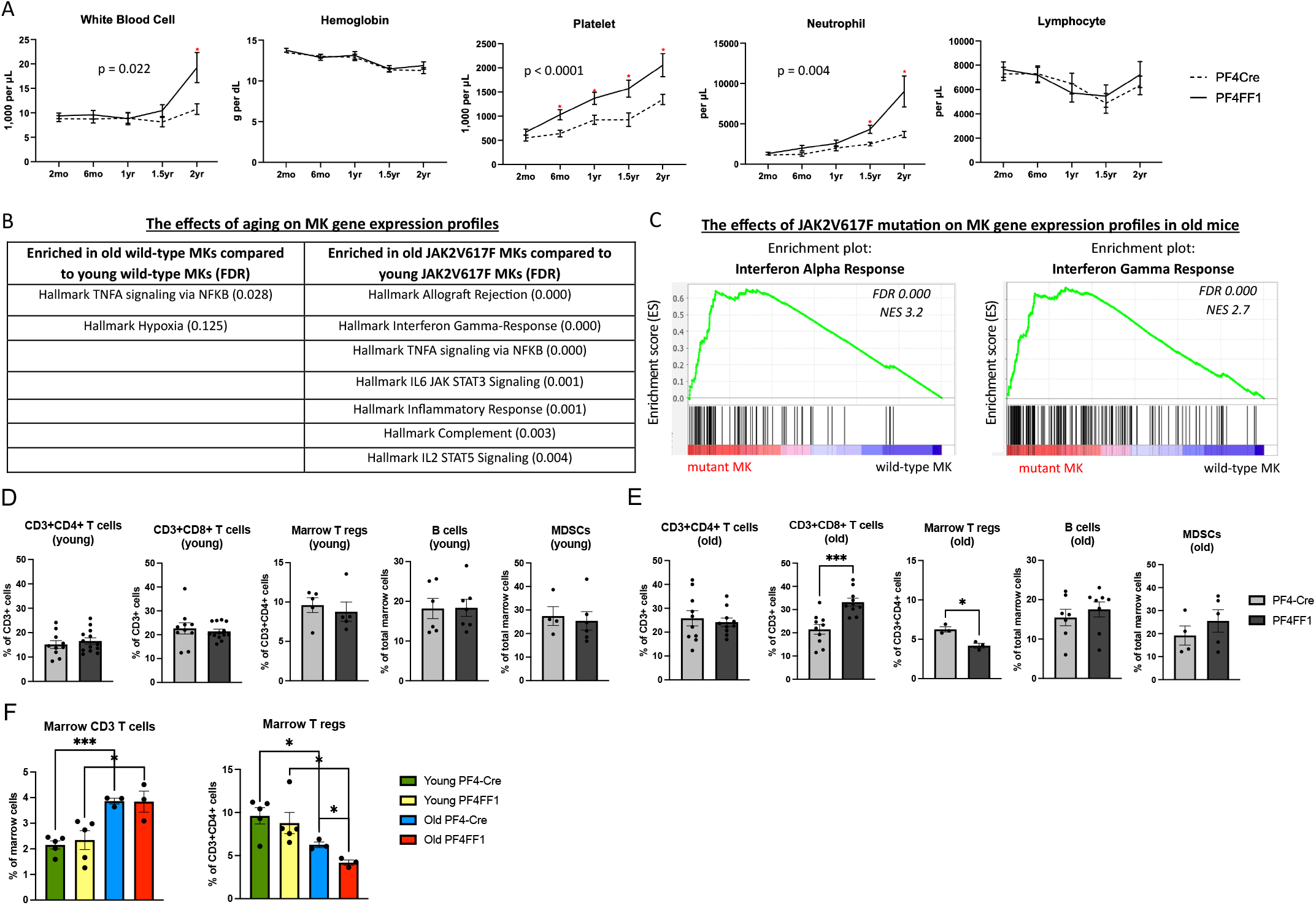
Inflammatory and immune activation signatures in JAK2V617F mutant MKs linked to skewed marrow T cell populations during aging of Pf4^+^FF1^+^ mice. (**A**) Peripheral blood cell counts of Pf4^+^FF1^+^ (black line) and Pf4-cre control mice (dotted line) (n=10-23 mice in each group). (**B**) Gene set enrichment analysis (GSEA) of differentially expressed genes in wild-type MKs (control mice; left) and JAK2V617F mutant MKs (Pf4^+^FF1^+^ mice; right) during aging. (**C**) Enriched interferon response pathways in JAK2V617F mutant MKs compared to wild-type MKs in old mice. (**D-E**) Frequency of bone marrow T cells (CD3^+^CD4^+^ helper T cells, CD3^+^CD8^+^ cytotoxic T cells, and CD3^+^CD4^+^CD25^+^FoxP3^+^ regulatory T cells), B cells (CD3^-^B220^+^), myeloid-derived suppressor cells or MDSCs (both CD11b^+^Ly6C^high^Ly6G^-^ M-MDSCs and CD11b^+^Ly6C^low^Ly6G^+^ PMN-MDSCs) in young (C) and old (D) Pf4-Cre control and Pf4^+^FF1^+^ mice (young: n=5-11 mice in each group; old: n=3-10 mice in each group). (**F**) Overall comparison of marrow CD3^+^ T cells and Treg cells in control and Pf4^+^FF1^+^ mice across aging (n=3-6 per group) (∗p < 0.05; ∗∗∗p < 0.001)

To investigate how JAK2V617F mutant MK function evolves during aging, we performed bulk RNA sequencing (RNAseq) on wild-type and JAK2V617F mutant CD41^+^ cells from young (6mo) and old (2yr) Pf4-cre control and Pf4^+^FF1^+^ mice^23^. While TNFα signaling via the NFκB pathway was upregulated in both wild-type and mutant MKs with aging, JAKV617F MKs exhibited a distinct gene expression profile of heightened inflammation and immune activation (Figure 1B). Notably, genes related to the Interferon alpha response and Interferon gamma response — two key regulators of the tumor microenvironment^29^ — were significantly upregulated in mutant MKs from aged Pf4^+^FF1^+^ mice compared to wild-type controls (Figure 1C).

Flow cytometry revealed a significant increase in CD3^+^CD8^+^ T cells and a concurrent decrease in T regulatory (Treg) cells in the marrow of aged Pf4^+^FF1^+^ mice compared to aged controls, changes not observed in young mice (Figure 1D-E). Overall, marrow CD3^+^ T cells increased with aging in both control and mutant mice, while Treg cells declined (Figure 1F). These findings underscore the impact of aging on the immune cell landscape within bone marrow, where mutant MKs exacerbate the increase in CD8^+^ T cells while depleting CD4^+^ Treg cells.

### Substantial changes in the marrow immune microenvironment of aging Pf4^+^FF1^+^ mice

To investigate how JAK2V617F mutant MKs alter the marrow immune microenvironment, we performed single-cell RNA-seq (scRNAseq) on unfractionated marrow cells from young (4mo) and aged (1yr) Pf4^+^FF1^+^ and control mice (n=1 per group). Following data integration, doublet removal and quality control (see Methods), 47,730 cells were analyzed from 4 mice (9,800 – 15,678 cells per mouse). Unsupervised clustering of all the data identified 29 clusters of marrow cells, visualized by uniform manifold approximation and projection (UMAP). Cell identity was manually assigned to clusters based on the differential expression of known marker genes, identifying distinct immune and hematopoietic populations including neutrophils, monocytes/macrophages, T/NK cells, B cells, dendritic cells, granulocyte-monocyte progenitors, MKs/HSPCs, eosinophil/basophil/mast cells (EBM), and erythroid cell populations (Figure 2A).

**Figure 2.**
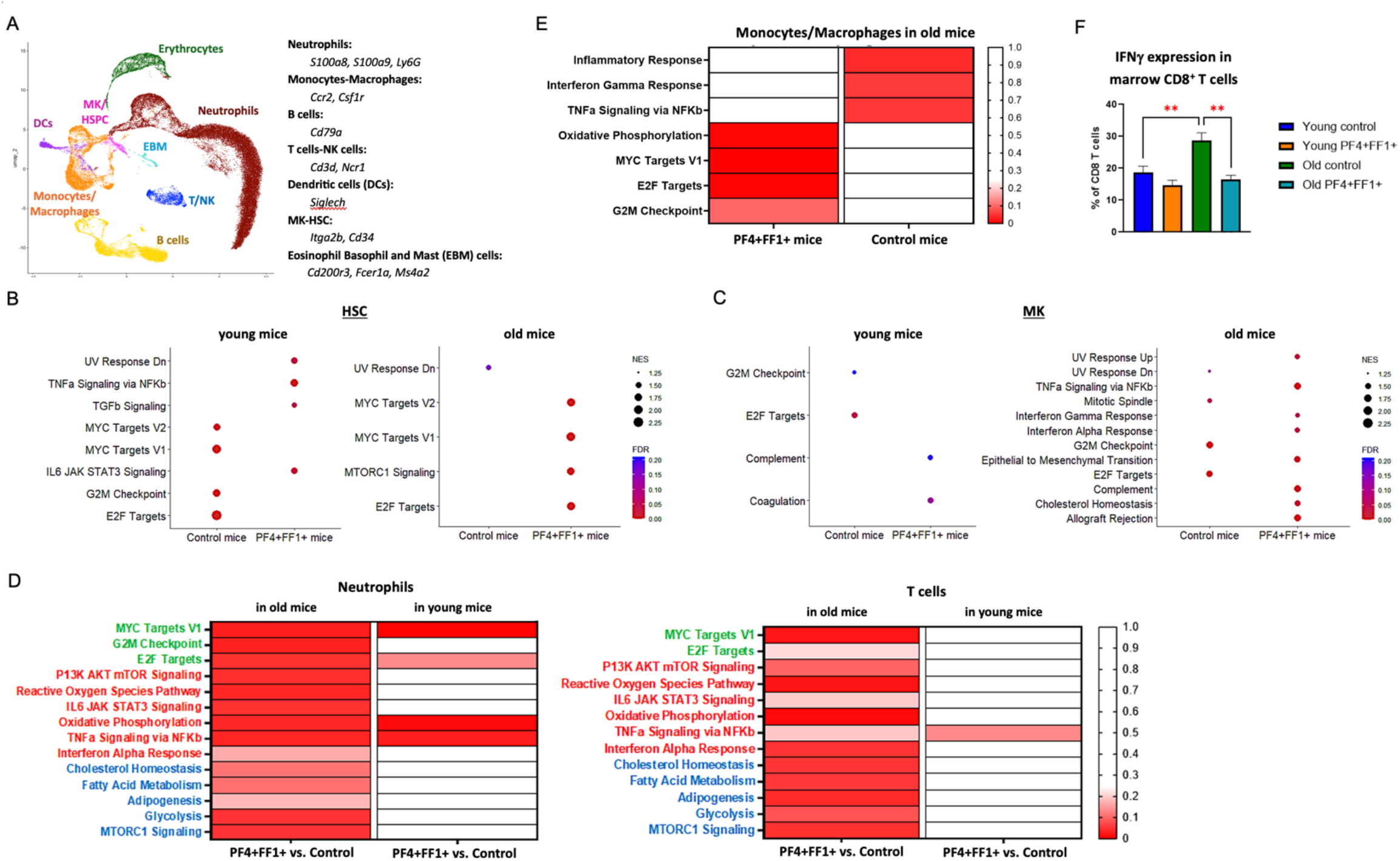
scRNAseq analysis of marrow cells from young and aged Pf4^+^FF1^+^ mice. (**A**) Uniform manifold approximation and projection (UMAP) plot of unfractionated marrow cells, with key markers used to infer cell identity listed. (**B-C**) GSEA analysis of HSCs (B) and MKs (C) from young (left) and old (right) Pf4^+^FF1^+^ and control mice. (**D**) Gene sets associated with cell proliferation (green), inflammation and immune activation (red), and metabolism (blue) in neutrophils and T cells from old (left) and young (right) Pf4^+^FF1^+^ mice compared to controls. (**E**) GSEA analysis of monocytes/macrophages in old Pf4^+^FF1^+^ (left) and control (right) mice. (**F**) Flow cytometry quantification of intracellular IFN-γ protein levels in bone marrow CD8^+^ T cells (n = 4-5 mice per group). (∗∗p < 0.01)

We performed an integrated cluster analysis of the MK/HSPC population. MKs were identified by their enriched expression of Itga2b, Pf4, Gp9, Mpl, and vWF, while HSCs were identified by their enriched expression of CD34, Flt3, and CD27. Consistent with previous findings of decreased HSC proliferation in young Pf4^+^FF1^+^ mice but increased activity in aged Pf4^+^FF1^+^ mice^23,30^, gene sets associated with proliferation (e.g., MYC targets, G2/M checkpoint, E2F targets) were significantly upregulated in HSCs from aged Pf4^+^FF1^+^ mice compared to age-matched controls, contrasting with their expression patterns in young mice (Figure 2B). Similarly, gene set enrichment analysis (GSEA) of MKs revealed that immune activation and inflammation pathways were significantly upregulated in aged Pf4^+^FF1^+^ mice, but not in young mice, reinforcing the idea that aging amplified mutant MK-driven inflammation (Figure 2C).

Beyond HSPCs and MKs, cell proliferation (e.g., MYC targets, G2/M checkpoint, E2F targets), inflammation and immune activation (e.g., PI3K AKT MTOR signaling, Reactive oxygen species pathway, IL6 JAK STAT3 signaling, Oxidative phosphorylation, TNFA signaling via NFKB, Interferon alpha response), and metabolic pathways (e.g., cholesterol homeostasis, fatty acid metabolism, glycolysis, MTORCS signaling) were highly upregulated in neutrophils and T cells in aged Pf4^+^FF1^+^ mice compared to age-matched controls, reflecting a pro-inflammatory marrow immune microenvironment (Figure 2D). In contrast, while cell proliferation pathways are significantly upregulated in monocytes and macrophages from aged Pf4^+^FF1^+^ mice, IL6/JAK/STAT3, TNFα/NFkB, interferon, and inflammatory response genes are notably downregulated, indicating a shift toward a less immune-active phenotype with aging (Figure 2E). In addition, despite the upregulation of proliferation and metabolism pathways in T cells, flow cytometry analysis revealed impaired CD8^+^ T cell cytotoxic function (measured by intracellular IFNγ levels) in aged Pf4^+^FF1^+^ mice (Figure 2F). These findings suggest that chronic inflammation induced by JAK2V617F mutant MKs skews T cell populations (Figure 1E), suppresses T cell function (Figure 2F), and promotes an immune-suppressive monocyte phenotype (Figure 2E).

### The modulatory functions of JAK2V617F mutant MKs in both innate and adaptive immunity

MHC class I molecules are expressed on all nucleated cells and are essential for presenting intracellular antigens to CD8^+^ T cells. IFNγ can upregulate MHC I gene expression through the activation of JAK-STAT signaling pathway^31^. Consistent with the upregulated interferon gamma response gene signature observed in JAK2V617F mutant MKs from aged Pf4^+^FF1^+^ mice compared to controls, flow cytometry analysis further validated the increased expression of MHC class I molecules on mutant MKs from aged Pf4^+^FF1^+^ mice (Figure 3A).

**Figure 3.**
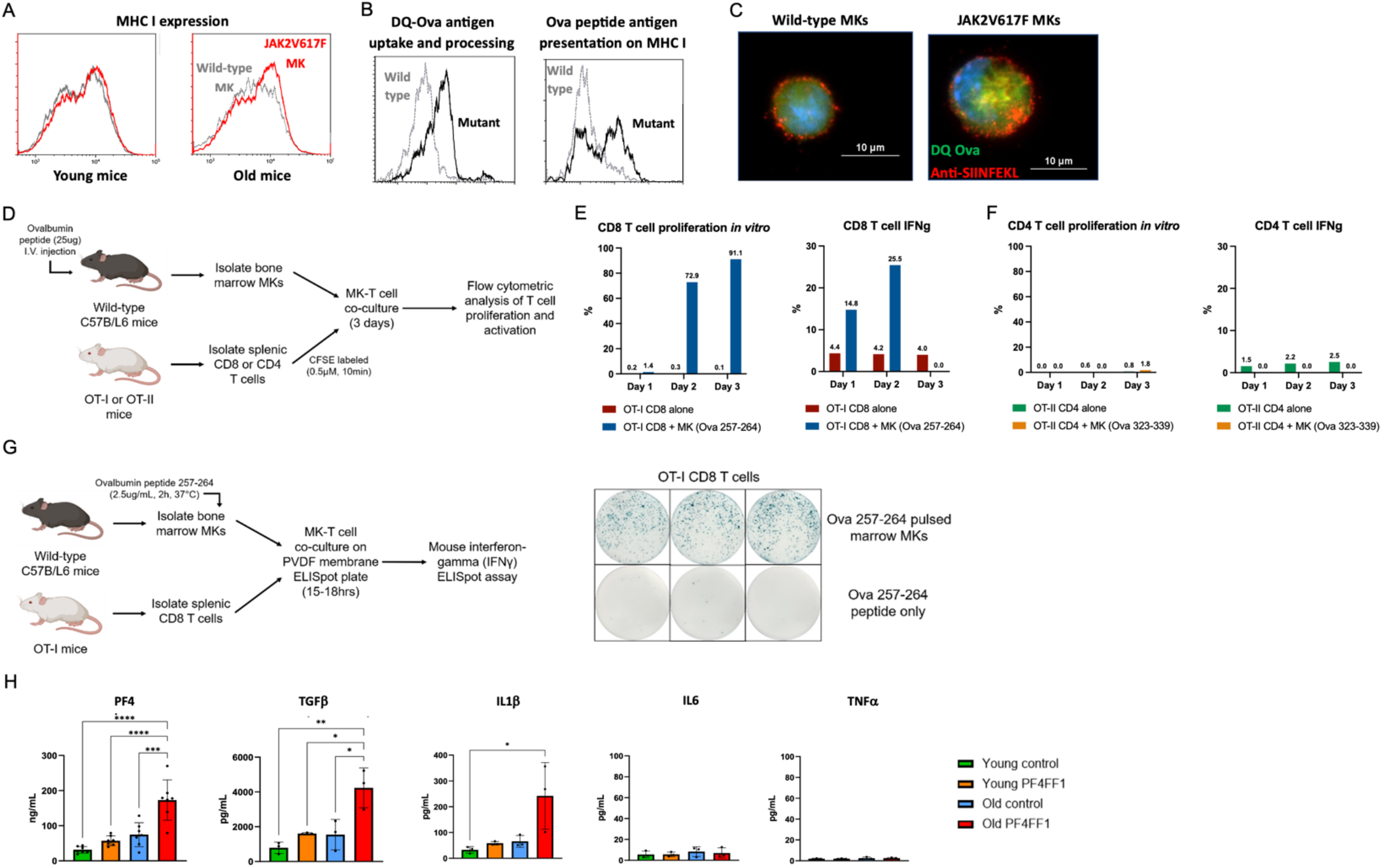
The immunomodulatory functions of JAK2V617F mutant MKs. (**A**) MHC I protein levels on wild-type and JAK2V617F MKs from young (left) and old (right) mice, measured by flow cytometry using an anti-mouse H-2D^b^ antibody (pooled data from two mice per group). (**B-C**) Representative flow cytometry analysis (B) and fluorescence microscope images (C) showing wild-type and JAK2V617F mutant MKs processing DQ-Ova (green fluorescence) and presenting Ova peptide antigen on MHC I molecules (red fluorescence). (**D**) *In vivo* experimental design using chicken ovalbumin antigen system to assess the impact of bone marrow MKs on T cell activation. (**E-F**): OT-I CD8^+^ T cell (E) and OT-II CD4^+^ T cell (F) proliferation (left) and IFNγ expression (right) following co-culture with marrow MKs from Ova-treated mice. (**G**) Experimental scheme (left) and IFNγ ELISpot assays (right) showing OT-I CD8^+^ T cell activation after co-culture with Ova257-284 pulsed MKs. (**H**) Protein levels of PF4, TGFβ, IL1β, IL6, and TNFα in marrow plasma from young and old Pf4^+^FF1^+^ mice and aged-matched controls (n=3-7 mice per group). (* p <0.05; ** p < 0.01; *** p < 0.001; **** p < 0.0001)

We then assessed the ability of JAK2V617F mutant MKs to process exogenous protein antigens^9,11^. Marrow MKs from aged wild-type and Pf4^+^FF1^+^ mice were cultured with 200ug/ml DQ-Ovalbumin (DQ-Ova) and cell fluorescence was measured by flow cytometry (Figure 3B) and fluorescence microscope (Figure 3C) 2hr later as a measure of Ova processing (DQ-Ova fluorescence is only activated upon cellular uptake and proteolytic cleavage). Antigen presentation on MHC I was evaluated using an anti-MHC class I-Ova antibody. JAK2V617F mutant MKs from aged Pf4^+^FF1^+^ mice showed enhanced Ova antigen uptake, processing, and presentation on MHC I compared to wild-type MKs from aged control mice.

To determine whether MKs interact with T cells in an antigen-specific manner, we used OT-I and OT-II transgenic mice, which recognize MHC I-restricted Ova257-264 and MHC II-restricted Ova323-339 peptides, respectively^32,33^. Wild-type mice were treated with a single dose of 25ug Ova257-264, Ova323-339, or PBS via intraperitoneal injection. One hour later, marrow MKs were isolated and co-cultured with naïve OT-I CD8^+^ or OT-II CD4^+^ T cells (Figure 3D). IFNγ expression and cell proliferation (by carboxyfluorescein succinimidyl ester, or CFSE label dilution) were assessed. Ova257-264-loaded marrow MKs significantly enhanced CD8+ T cell proliferation and transiently increased IFNγ production, peaking at Day 2 (Figure 3E). In contrast, Ova323-339-loaded marrow MKs had no effect on CD4^+^ T cells (Figure 3F). To confirm that Ova257-264-stimulated MKs directly activate CD8^+^ T cells, wild-type marrow MKs were pulsed with Ova257-264 *in vitro* (2.5ug/ml, 2 hours at 37°C) before co-culture with OT-I splenic CD8^+^ T cells. A mouse IFNγ Enzyme-linked immunospot (ELISpot) assay revealed a significant increase in IFNγ-secreting CD8^+^ T cells, consistent with *in vivo* findings (Figure 3G). We observed no significant differences in T cell proliferation and activation between wild-type and JAK2V617F mutant MKs (data not shown), indicating that both can interact with T cells in an antigen-specific manner.

An important function of innate immunity is cytokine production to modulate immune responses. ELISA analysis of platelet-free marrow plasma (collected by flushing each murine femur with 0.5ml PBS) showed significantly elevated PF4 (Platelet factor 4), TGFβ (Transforming growth factor beta), and IL1β (Interleukin-1 beta) in aged Pf4^+^FF1^+^ mice compared to controls, with no differences in young mice (Figure 3H). MKs are the primary source of PF4 in the marrow^4^, and its plasma levels are elevated in many MPN patients^34,35^. PF4 is involved in immune cell recruitment^36^, monocyte differentiation^37,38^, and CD8^+^ T cell inhibition^37,39^. Notably, in aged Pf4^+^FF1^+^ mice marrow plasma, PF4 (170ng/ml) was markedly higher than TGFβ (4ng/ml) and IL1β (0.2ng/ml), suggesting that PF4 upregulation may play a key role in JAK2V617F mutant MK-induced immune dysfunction.

Taken together, JAK2V617F mutant MKs upregulate inflammatory and immune regulatory genes, produce pro-inflammatory cytokines, present antigens, and modulate T cell functions, thereby driving the development of an inflammatory hematopoietic microenvironment. Given that chronic inflammation is a hallmark of the MPN hematopoietic niche and promotes the expansion of JAK2V617F mutant cells over their wild-type counterparts^40^, we propose that the inflammatory marrow microenvironment, shaped by the immune modulatory activities of JAK2V617F mutant MKs, plays a critical role in MPN disease progression.

### Increased inflammatory and immunomodulatory gene expression in MKs is associated with MPN disease progression in a murine model with co-existing wild-type and JAK2V617F mutant cells

To test whether this heightened immune modulatory MK signature is linked to MPN disease progression, we generated chimeric murine models harboring both wild-type and JAK2V617F mutant hematopoietic cells at varying ratios to model MPN disease progression. We used the Tie2-cre^+^FF1^+^ murine model we previously established in which human JAK2V617F is expressed in all hematopoietic cells^41-43^. In brief, lethally irradiated wild-type mice (CD45.1) were transplanted with both wild-type marrow cells (isolated from CD45.1 wild-type mice) and JAK2V617F mutant marrow cells (isolated from CD45.2 Tie2-cre^+^FF1^+^ mice) mixed at ratios of 100:0, 90:10, 10:90. These chimeric mice exhibited a “dose-dependent” MPN phenotype, with those receiving a higher proportion (90%) mutant donor cells developing leukocytosis, thrombocytosis, and an increased marrow Lin^-^ cKit^+^Sca1^+^ (LSKs) and Lin^-^cKit^+^Sca1^+^CD150^+^CD48^-^ (HSCs) 16 weeks post-transplantation, while those receiving a lower proportion (10%) of mutant donor cells remained asymptomatic throughout the 4-month follow-up period (Figure 4A).

**Figure 4.**
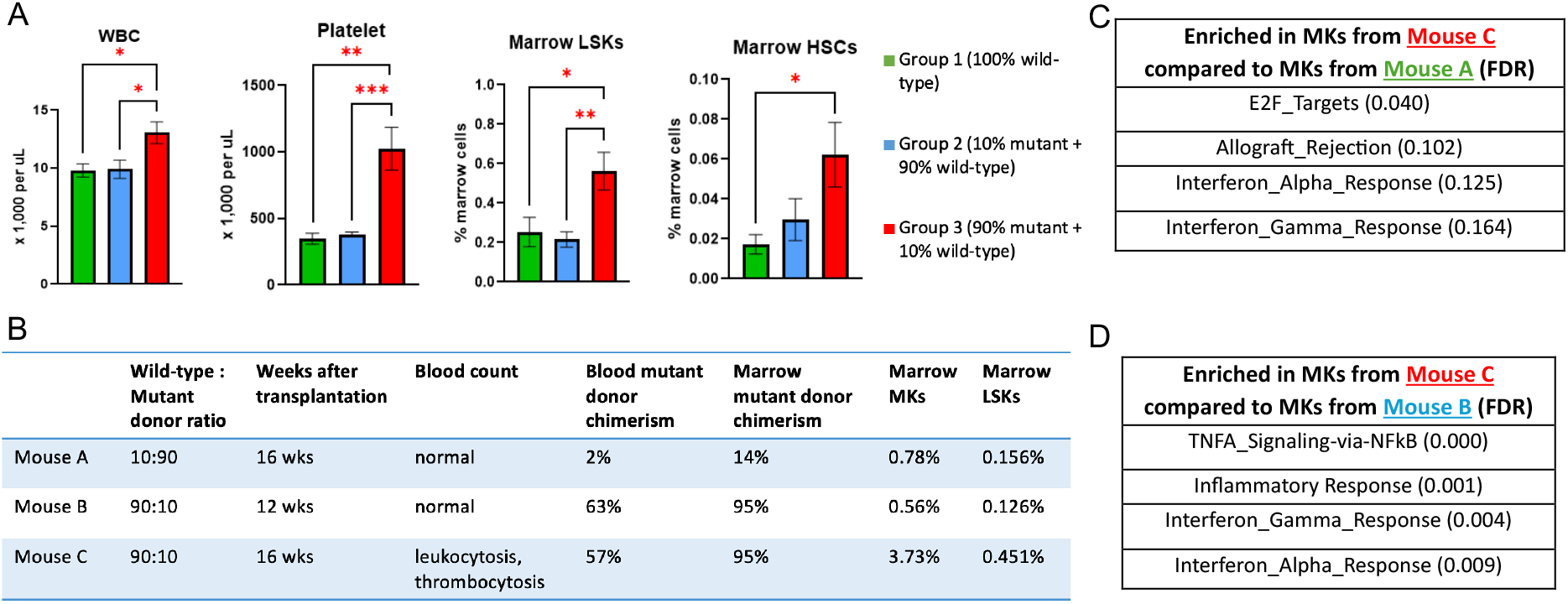
Augmented inflammatory and immunomodulatory gene expression in MKs is associated with MPN disease progression. (**A**) Peripheral blood WBC and platelet counts, marrow LSK and HSC numbers of the chimeric mice with varying mutant cell burdens at 16wks post transplantation (n=6-8 mice in each group). (**B**) Mice selected for scRNAseq analysis. (**C**) Top gene sets significantly enriched in mouse C MKs compared to mouse A MKs. (**D**) Top gene sets significantly enriched in mouse C MKs compared to mouse B MKs. (∗p < 0.05; ∗∗p < 0.01; ∗∗∗p < 0.001)

We conducted scRNAseq analysis on unfractionated marrow cells from three chimeric mice (A, B, and C), characterized by varying mutant cell burdens and distinct hematologic phenotypes at different time points post-transplantation (Figure 4B). Following quality filtering of the sequence data, a total of 15,403 cells from three mice were included in the analysis. MKs were identified by the expression of established marker genes, including Itga2b, Pf4, Gp9, vWF, and MPL. Comparative analysis of gene expression profiles revealed significant enrichment of cell proliferation, inflammation, and immune activation gene sets in MKs from a high mutant burden MPN mouse (Mouse C) compared to MKs from a low mutant burden, non-MPN mouse (Mouse A) (Figure 4C). Similarly, MKs from Mouse C showed the same enrichment when compared to MKs from a comparable mutant burden, non-MPN mouse (Mouse B) (Figure 4D). These findings suggest that heightened immunomodulatory signatures in MKs are linked to mutant cell expansion and MPN progression (mouse C vs. mouse A) and that the immunomodulatory functions of mutant MKs are not solely attributable to the presence of the JAK2V617F mutation alone (mouse C vs. mouse B).

### Elevated LINE-1 transcription in bone marrow MKs in MPN murine models

Half of the human genome consists of transposable elements, with LINE-1 (long-interspersed element-1) being the only protein-coding transposon that remains active in humans^44^. While LINE-1 reverse transcriptase activity has been detected in human and mouse platelets^45^, little is known about its expression in MKs. Since LINE-1 activation is linked to DNA replication and cell cycling^46^, and MKs undergo endomitosis — a unique form of cell cycling during which MKs undergo multiple rounds of DNA synthesis without cell division^47^ — it is likely that LINE-1 activity increases in MKs with aging.

Full-length LINE-1 elements in mice span ∼7kb, containing a 5’ UTR, two open reading frames ORF1 and ORF2, and a 3’ UTR with a poly(A) tail. RT-qPCR using multiple primer pairs targeting active LINE-1 families^48^ revealed a significant increase in 5’UTR, ORF1, ORF2, and 3’UTR transcripts in mutant MKs from aged mice, but not young mice (Figure 5A). Treating marrow MKs from aged control and Pf4^+^FF1^+^ mice with lamivudine (25uM for 24hrs), a reverse transcriptase inhibitor used to suppress HIV replication, significantly reduced LINE-1 transcripts in mutant MKs (Figure 5B). This suggests that reverse transcription plays a role in the elevated LINE-1 RNA levels in JAK2V617F MKs. Given that LINE-1 activation is implicated in aging-related inflammation^48-51^, these findings suggest that elevated LINE-1 expression in JAK2V617F mutant MKs may activate the innate immune response and contribute to an inflammatory marrow microenvironment.

**Figure 5.**
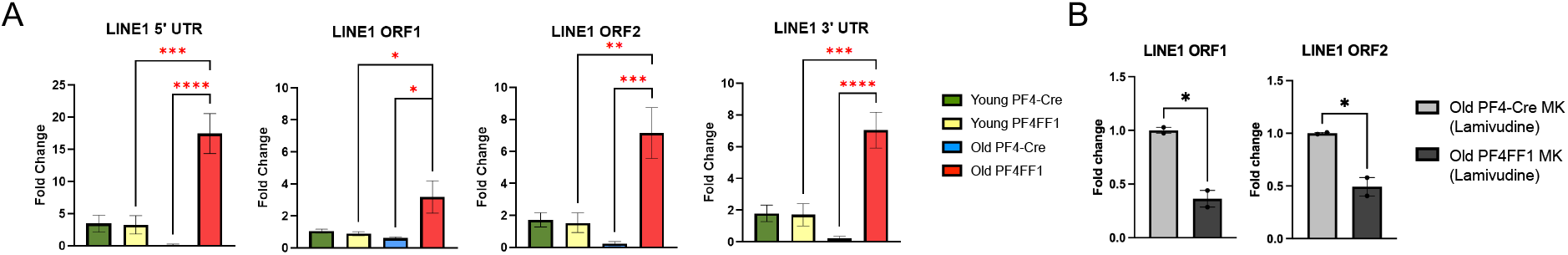
Elevated LINE-1 expression in JAK2V617F mutant MKs during aging. (**A**) LINE-1 RNA levels were evaluated using primers specific to the 5’ UTR, ORF1, ORF2, and 3’ UTR. The transcript levels are shown as fold changes relative to their expression levels in MKs from young control mice. (n=3-4 mice in each group). (**B**) ORF1 and ORF2 RNA levels in control and JAK2V617F mutant marrow MKs following lamivudine treatment. (∗p < 0.05; ∗∗p < 0.01; ∗∗∗p < 0.001)

### Increased endocytosis and antigen processing of exogenous ovalbumin by iPS-derived JAK2V617F mutant human MKs compared to wild-type MKs

Studying human disease could be challenging due to the limited availability of relevant cells and tissues. Primary MKs from patients are typically a heterogeneous mix of normal and diseased cells, making their functional analysis complicated due to the genetic diversity. Reprogramming of somatic cells to an embryonic stem cell-like state (iPS cells) has made it possible to create disease models that retain patient-specific genetic alterations. For this study, we obtained a JAK2V617F homozygous mutant iPS cell line derived from a MPN patient (PVB1.4) and a JAK2 wild-type iPS cell line derived from a healthy donor (BC1)^52^. Their genotypes were confirmed by a nested allele-specific PCR assay^53^ (Figure 6A-B). As expected, human JAK2 gene expression is significantly higher in PVB1.4 cells than in BC1 cells (Figure 6C). These iPS cells can be differentiated into CD34^+^CD45^+^ hematopoietic progenitors using the spin-embryoid body method, harvested after 2-3 weeks, and further differentiated into CD41^+^CD42^+^ mature MKs^54-56^ (Figure 6D-E).

**Figure 6.**
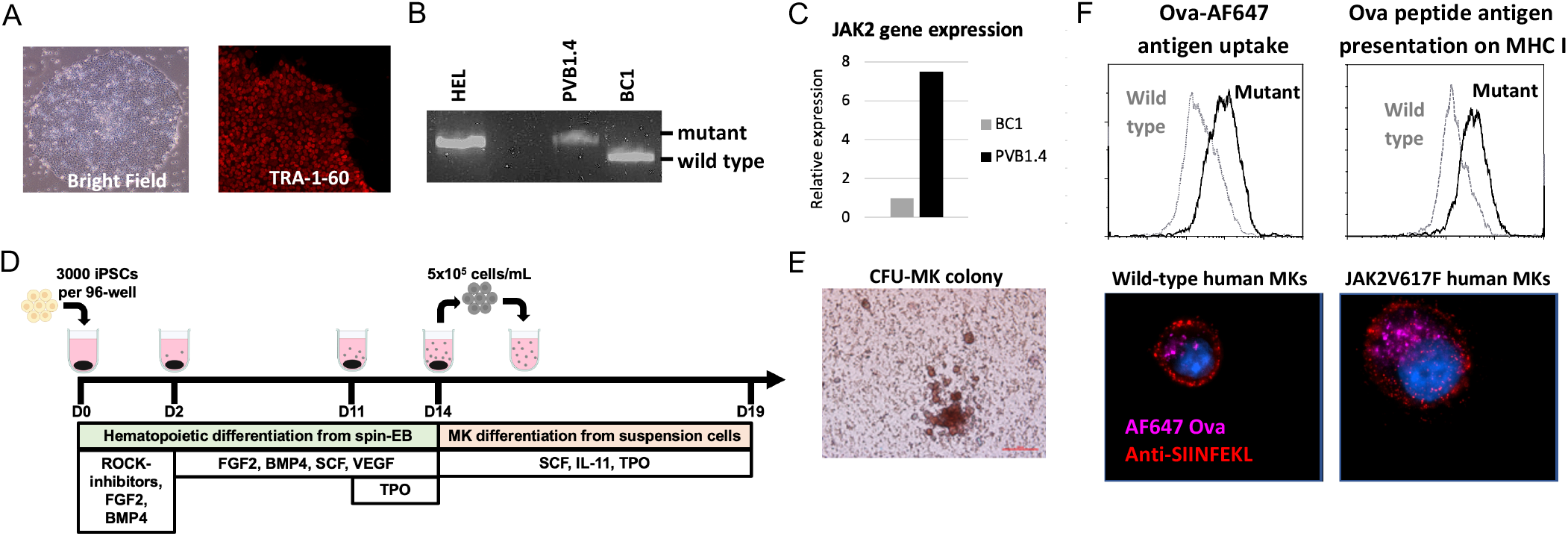
iPS cell line model for studying human MK functions. (**A**) Immunofluorescence staining of iPS colonies showing expression of the undifferentiated cell markers TRA1-60. (**B**) Detection of wild-type JAK2 allele (229bp) and JAK2V617F mutant allele (279bp) by nested allele-specific PCR. HEL: human erythroleukemia cell line (JAK2V617F mutant, positive control). (**C**) JAK2 gene expression in BC1 and PVB1.4 iPS cells measured by real-time qPCR. Gene expression is shown as fold-change relative to BC1, which was set as 1. (**D**) Schematic presentation of the hematopoietic and megakaryocytic differentiation strategy for human iPS cells. (**E**) Representative CFU-MK colony formation assay of iPS-derived CD41^+^ MKs. (**F**) Representative flow cytometry analysis (top) and fluorescence microscope images (bottom) of wild-type and JAK2V617F mutant iPS-derived human MKs uptake Ova (purple fluorescence) and present Ova peptide antigen on MHC I molecules (red fluorescence). Magnification: 1000X.

**Figure 7.**
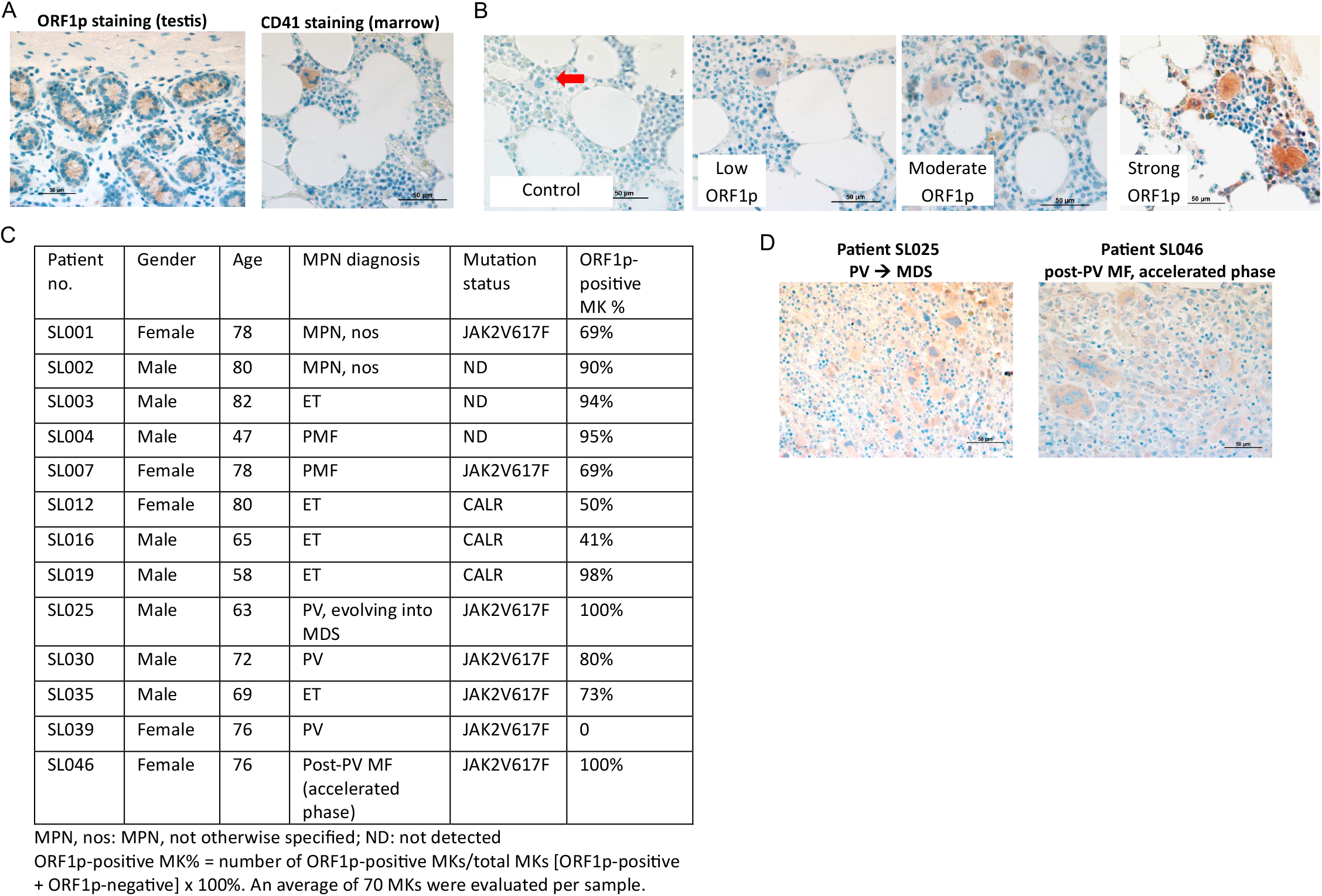
LINE-1 ORF1p protein expression in marrow MKs from MPN patients. (**A**) Representative images of ORF1p-positive staining in a histologically normal adult testis (left) and CD41 staining in normal bone marrow (right). (**B**) ORF1p protein expression in marrow biopsies from a representative control patient and three MPN patients with low, moderate, and strong ORF1p expression. (**C**) Summary of gender, age, MPN diagnosis, and ORF1p-positive MK (%) in the thirteen MPN patients analyzed. (**D**) Diffuse ORF1p expression in the marrow of two MPN patients with disease progression. All images were captured at 400x magnification.

To validate murine findings in human cells, we assessed the antigen-processing ability of JAK2V617F mutant human MKs. Wild-type (BC1-derived) or JAK2V617F mutant (PVB1.4-derived) MKs were cultured with 200ug/ml Ovalbumin-Alexa Fluor 647 for 2 hours at 37ºC. Similar to murine MKs (Figure 3B-C), JAK2V617F mutant human MKs exhibit increased Ova uptake and presentation on MHC class I compared to wild-type MKs (Figure 6F).

### LINE-1 ORF1p protein expression in marrow MKs of MPN patients

Finally, we assessed LINE-1 expression in marrow biopsies from MPN patients (n=13, average age 71yr) and age-matched controls from orthopedic surgery patients (4 hip marrow samples and 1 rib marrow samples; average age 71yr) using immunohistochemistry (IHC) for CD41 and ORF1p. ORF1p staining was validated in adult testis tissue, known to low ORF1p expression^57^, and MKs were identified based on morphology and CD41 positivity (Figure 7A). No ORF1p protein was detected in any of the five control marrow samples. In contrast, ORF1p expression was observed in marrow MKs in 12 out of 13 MPN patients, with varying levels of positivity (Figure 7B). Within individual MPN patients, both ORF1p-positive and ORF1p-negative MKs co-existed, with 40-100% MKs expressing ORF1p in affected patients (Figure 7C). Notably, while ORF1p expression was largely restricted to MKs, diffuse ORF1p expression in non-MK marrow cells was observed in two patients with advanced disease — one transitioning from PV to myelodysplastic syndrome (MDS) (patient #SL025) and another with accelerated-phase post-PV myelofibrosis (MF) (patient #SL046) (Figure 7D). These findings suggest that LINE-1 activation is a hallmark of MPN marrow MKs, with its expression increasing in advanced disease stages. The presence of ORF1p in non-MK marrow cells in progressive MPN cases further raises the possibility that LINE-1 activation extends beyond MKs during disease evolution, potentially contributing to neoplastic clonal expansion.

## Discussion

MKs, long recognized for their role in platelet production, are increasingly understood as key regulators of the hematopoietic and immune microenvironment. By producing inflammatory cytokines and immune mediators, MKs actively participate in pathogen surveillance and immune responses in tissue microenvironment^7-12,58^. However, how diseased MKs alter these functions to influence HSC behavior in neoplastic hematopoiesis remains unclear. Our findings establish JAK2V617F mutant MKs as potent immunomodulators, capable of reprogramming the marrow immune microenvironment and driving disease progression in MPNs. Compared to wild-type MKs, mutant MKs exhibit heightened inflammation and innate immune activation, including increased antigen presentation, elevated pro-inflammatory cytokines (PF4, TGFβ, IL1β), skewed T cell populations, and impaired T cell functions in the JAK2V617F-bearing MK niche. These changes are further amplified by aging, a major risk factor for MPN progression. To determine the impact of MK-driven immune dysregulation on MPN evolution, we developed a chimeric murine model with co-existing wild-type and JAK2V617F mutant HSCs, mimicking the competitive dynamics observed in MPN patients. Our results reveal that enhanced MK immunomodulatory function is linked to mutant cell expansion and disease progression, underscoring MKs’ role as key drivers of clonal hematopoiesis. These findings were further validated in human cells using an MPN patient-derived iPS cell line model, which allowed us to confirm increased antigen processing and presentation in JAK2V617F mutant MKs compared to wild-type MKs in a genetically controlled system.

Approximately half of the human genome consists of transposable elements, with LINE-1 being the only protein-coding transposon still active in humans^44^. As LINE-1 replicates itself using RNA intermediates via retrotransposition, LINE-1-derived endogenous nucleic acids can act as danger signals to trigger the innate immune response via pattern recognition receptors, leading to immune cell activation and the production of inflammatory cytokines^59,60^. In recent years, it has been increasingly recognized that LINE-1 activation is a universal hallmark of aging and a key driver of innate immune activation^48-51^. LINE-1 retrotransposition activity peaks during cell cycling and DNA replication^46^. MKs, which undergo endomitosis and accumulate large amounts of DNA and RNA, are uniquely positioned to be susceptible to LINE-1 reactivation. However, the role of LINE-1 in MKs has remained largely unexplored. Here, we show that the LINE-1 transcripts are significantly upregulated in aging JAK2V617F mutant MKs in an MPN murine model, a finding corroborated by the expression of LINE-1–encoded protein ORF1p marrow MKs from 13 out of 14 MPN patients, but absent in age-matched orthopedic surgery controls. These findings suggest that LINE-1 activation in MKs serve as a key inflammatory trigger in the MPN marrow niche. Furthermore, our data demonstrate that treating JAK2V617F mutant MKs with a reverse transcriptase inhibitor significantly reduces LINE-1 expression, suggesting that reverse transcription plays an important role in the elevated LINE-1 RNA levels in JAK2V617F MKs and supporting the idea that LINE-1– derived nucleic acids trigger a type I interferon response, fueling chronic inflammation and clonal expansion.

While chronic inflammation is a well-established driver of HSC stress and aging-associated hematological disorders, the specific niche cells responsible for inflammatory cytokine production remain poorly defined. Our findings suggest that MKs are central regulators of niche inflammation in MPNs, influencing both innate and adaptive immunity to create a marrow microenvironment that suppresses wild-type hematopoiesis while promoting neoplastic HSC expansion. Given that aging-related decline in normal hematopoiesis contributes to clonal hematopoiesis by increasing the relative fitness of mutant HSCs, our study identifies mutant MKs as a key link between aging, inflammation, and neoplastic progression.

The translational significance of these findings extends beyond MPNs. Abnormal megakaryopoiesis is a common feature of many hematologic malignancies^13-18^, and aging-associated clonal hematopoiesis is increasingly recognized as a risk factor for cardiovascular disease, immune dysregulation, and cancer^61-64^. Our study highlights the broader role of MK-driven inflammation in shaping the marrow and systemic immune landscape, with implications for developing novel therapeutic strategies targeting inflammatory and immune pathways activated in mutant MKs. Targeting JAK-STAT signaling, MHC class I presentation, cytokine production, or LINE-1 reactivation could provide new approaches to suppress inflammation and restore immune balance in MPNs and other clonal hematopoietic disorders. While this study establishes MKs as central players in marrow inflammation and immune modulation, several key questions remain. How does JAK2V617F promote LINE-1 activation in MKs? Does MK-driven inflammation cause or accelerate MPN disease progression, or is it a consequence of clonal dominance? Answering these questions will be crucial for developing targeted therapies aimed at disrupting the vicious cycle of mutant MK-driven inflammation, immune dysfunction, and clonal expansion.

## Acknowledgements

This research was supported by the VA Merit Award BX003947 and BX005584 (H.Z.) and NIH R01 HL134970 and R01 CA266294 (H.Z.). We thank Dr. Kathleen Burns (Dana-Farber Cancer Institute, Harvard Medical School) for her valuable insights and discussions on the LINE-1 research.

## Author Contributions

S.L. performed transgenic murine model experiments and data analysis; X.Y. conducted patient sample studies and assisted with murine studies; K.M. contributed to transgenic murine model studies and scRNAseq data analysis; T. Ahmed provided patient samples; L. Zheng provided scientific and technical support for immunology studies; H. Zhan designed and supervised the experiments, analyzed data, interpreted results, and wrote the manuscript. All authors read and approved the manuscript.

