## Supplemental data for "The Immune-Modulatory Function of Megakaryocytes in the Hematopoietic Niche of Myeloproliferative Neoplasms"

**Materials and Methods**

***Complete blood counts***

Peripheral blood was obtained from the facial vein via submandibular bleeding and collected in EDTA tubes. The samples were then analyzed using a Vetscan Hm5 Hematology Analyzer (Abaxis) to determine the blood count.

***Marrow cell isolation***

Murine femurs and tibias were first harvested and cleaned thoroughly. Marrow cells were flushed into PBS with 2% fetal bovine serum using a 25G needle and syringe. Remaining bones were crushed with a mortar and pestle followed by enzymatic digestion with DNase I (25U/ml) and Collagenase D (1mg/ml) at 37 °C for 20 min under gentle rocking. Both flushed and digested marrow tissues were thoroughly dissociated by gentle and repeated mixing using a 10ml pipette to facilitate dissociation of cellular aggregates. Resulting cell suspensions were then filtered through a 70uM cell strainer.

***Flow cytometry***

Cells were counted and stained with antibodies by incubating cell suspensions at 4°C for 30 minutes in the dark. Samples were analyzed using a Cytoflex flow cytometer (Beckman Coulter, CA, USA) or sorted using a FACSAria III (BD biosciences, CA, USA). CD3 (clone 17A2, Biolegned), CD4 (clone GK1.5, Biolegend), CD8 (clone 53-6.7, Biolegend), CD25 (clone 3C7, Biolegend), FoxP3 (clone MF-14, Biolegend), CD45R/B220 (clone RA3-6B2, Biolegend), CD11b (clone M1/70, Biolegend), Ly6C (clone HK1.4, Biolegend), Ly6G (clone 1A8, Biolegend), CD41 (clone MWReg30, Biolegend), H-2D$^b$ (clone KH95, Biolegned), IFN$\gamma$ (clone XMG1.2, Biolegend), LINE1 ORF1p (clone ERP21844-108, Abcam), and Alexa Fluor 488 goat anti-rabbit IgG (A11008, Thermo Fisher, Waltham, MA) antibodies were used. Flow cytometry data were analyzed using Kaluza (Beckman Coulter).

***Transcriptome analysis using bulk RNA sequencing***

Wild-type and JAK2V617F mutant marrow CD41$^+$ cells were isolated from 3-4mo old young PF4-cre control mice (n=3) and PF4FF1 mice (n=3), and 2yr old Pf4-cre control mice (n=3) and Pf4$^+$FF1$^+$ mice (n=4), using a PE-conjugated anti-mouse CD41 antibody (Biolegend) and EasySep Mouse PE Positive Selection Kit II (Stem Cell Technologies). ~95% of these isolated cells express CD41 as measured by flow cytometry analysis. Total RNA was extracted using the RNeasy mini kit (Qiagen, Hilden, Germany). RNA integrity and quantitation were assessed using the RNA Nano 6000 Assay Kit of the Bioanalyzer 2100 system (Agilent Technologies, CA, USA). Bulk RNA sequencing was performed as previously described (Novogene, Durham, NC, USA)[1]. Differential gene expression analysis of MKs from young PF4-cre control mice (n=3), young PF4FF1 mice (n=3), old PF4-cre control mice (n=3), and old PF4FF1 mice (n=4), was performed using the DESeq R package (1.18.0). Genes with a *P*-value < 0.05 found by DESeq were assigned as differentially expressed.

***Single cell RNA sequencing***

Murine femurs and tibias were harvested and cleaned thoroughly. The bones were finely cut in PBS with 0.04% BSA. Cell suspensions were thoroughly homogenized by gentle and repeated mixing using a P1000 micropipette to facilitate dissociation of cellular aggregates. Resulting cell suspensions were then filtered through a 70uM cell strainer. Red blood cells were lysed twice using Red Blood Cell Lysing Buffer Hybri-Max (Sigma R7757). Final single bone marrow cells were washed and prepared in PBS with 0.04% BSA at a cell density of approximately $10^6$ cells per 1 mL. Freshly isolated cell suspensions were immediately processed at the Single Cell Core Facility (Stony Brook University and Northport VA Medical Center) to generate single cell RNA sequencing libraries using Chromium Next GEM Single Cell 5′ Reagent Kit v2 (10x Genomics, Pleasanton, CA) according to the manufacturer's instructions. Briefly, cells, gel beads, and partitioning oil were loaded into a Chromium Next GEM Chip G, for a target recovery of 10,000 cells per sample. The chip was loaded into a Chromium Controller (10x Genomics) to generate Gel Bead-in-Emulsions (GEMs). After reverse transcription using a GEM–RT incubation protocol on a PCR cycler, the GEMs were broken using Recovery Agent (10x Genomics), and the cDNA was purified with DynaBeads MyOne Silane beads (10x Genomics), amplified by PCR for 13 cycles, and further purified with SPRIselect reagent (Beckman Coulter, Brea, CA, USA). The DNA concentration and amplicon size were measured using a TapeStation 4200 with the D5000 high sensitivity ScreenTape system (Agilent, Santa Clara, CA, USA). After fragmentation, end-repair, A-tailing, and size selection, the cDNA was ligated with adaptors and purified again with SPRISselect reagents, according to 10x Genomics instructions. Libraries were amplified by PCR with sample index oligonucleotide primers from the Chromium i7 Multiplex Kit (10x Genomics), for a total of 10 or 11 cycles, depending on the initial cDNA yield. The final PCR products were subjected to double-sided size selection with SPRISselect reagents. Library cDNA concentration and size was determined on the Agilent TapeStation 4200. cDNA libraries were sequenced using an NovaSeq XPlus, PE150 with the following sequencing settings: Read 1 – 151bp; i7 index – 10bp; Read 2 – 151bp (Novogene, Durham, NC, USA).

***Single cell RNA sequencing data analysis***

Raw sequencing data FASTQ files were aligned to the transcriptome assembly mm10-2020-A. Counts were obtained using the CellRanger count function, then analyzed using the Seurat 4.3.0 (R Foundation for Statistical Computing, Vienna, Austria). Correction of the gene expression matrix for ambient RNA contamination was done using the SoupX R package. Determination of doublets for exclusion was done using the DoubletFinder R package. Cells were filtered to have at least 200 unique genes and less than 10% mitochondrial genes, as well as a log-normalized genes per UMI ratio of > 0.8. Data were normalized using the NormalizeData function using default parameters, and the top variable genes were identified using the FindVariableFeatures function with parameters of selection.method = “vst” and features = 3,000. After scaling the data using the ScaleData function, the most variable genes were used for principal component analysis (PCA) dimensionality reduction, using PCs 1:20 as determined by an Elbow Plot. Data were then integrated and the resulting clusters were visualized in a 2D Uniform Manifold and Approximation Projection (UMAP). Markers for identification were determined with a combination of literature review for markers, the use of marker resources such as CellMarker2.0 and Cell Taxonomy, information from companies such as Biolegend and R&D, and visualization of these markers in UMAP and violin plots. Differential expression data between clusters was retrieved using the FindMarkers function. The FindMarkers function was also used to retrieve differential expression data for between samples, ranked by p adjusted value.

***In vitro murine MK antigen uptake, processing, and presentation***

$CD41^+$ murine MKs were isolated from bone marrow using the EasySep Positive Selection Kit (StemCell Technologies, Cat. 17668) following the manufacturer’s protocol. Isolated MKs were cultured overnight (~16 hours) in Serum-Free Expansion Media (SFEM, StemCell Technologies) supplemented with 25ng/mL recombinant mouse SCF and 25ng/mL recombinant mouse TPO at a density of $2x10^5$ MKs/ml in untreated round-bottom 96-well plates. Cells were then incubated with DQ-Ovalbumin (DQ-Ova, Thermo Fisher, Cat. D12053) at 200 ug/mL for 1 hour at 37°C. After incubation, cells were washed three times using cold PBS with 2% FBS. Cells were then stained with a monoclonal antibody targeting the OVA257-264 (SIINFEKL) peptide bound to H2-Kb of MHC class I (Invitrogen, Cat. 14574381) at 0.125 µg/test for 30 minutes at 4°C, followed by washing and staining with an Alexa Fluor 555-conjugated goat anti-mouse secondary antibody (Invitrogen, A-21422, 1:1000 dilution) for 30 minutes at 4°C. Cells were then washed, and fluorescence was analyzed by flow cytometry to assess antigen processing (DQ-Ova fluorescence is activated upon proteolytic cleavage) and antigen presentation (detected by binding of the anti-SIINFEKLE antibody to MHC I). For immunofluorescence imaging, cells were fixed with 4% PFA for 10 minutes at room temperature, washed, and stained with DAPI (2ug/mL) for 10 minutes at room temperature. Cells were then washed, cytospun onto a glass slide, and imaged using a Nikon Ts2R-FL microscope.

A similar protocol was applied to human $CD41^+$ MKs differentiated from iPS cell lines. In brief, iPS-differentiated MKs were cultured in SFEM containing 50ng/mL recombinant human SCF and 50ng/mL recombinant human TPO. Cells were incubated with Ovalbumin conjugated with an AF647 for 2 hours at 37ºC to assess antigen uptake. Antigen processing and presentation were evaluated by staining with the OVA257-264 (SIINFEKL)-H2-Kb MHC class I antibody, followed by flow cytometry analysis.

***Measurement of marrow plasma cytokines***

Bone marrow plasma was collected by flushing 0.5mL of PBS into the marrow cavity of a single femur. The fluid was collected and centrifuged at 2,000 g for 15min to remove cell debris. For measurement of Platelet factor 4 (PF4) and Transforming growth factor beta (TGFβ) levels, samples underwent an additional centrifugation step to ensure complete platelet removal. The resulting marrow plasma was aliquoted and stored at -80℃ until further analysis. Cytokines levels were measured using ELISA kits following the manufacturer’s instructions: PF4 (R&D Systems, Cat. MCX400), TGFβ (R&D Systems, Cat. DB100C), IL1β (R&D Systems, Cat. MLB00C), TNFα (R&D Systems, Cat. MTA00B-1), IL6 (R&D Systems, Cat. M6000B-1) ELISA kit according. All samples were analyzed in duplicate.

***Real-time quantitative polymerase chain reaction***

The Azure Cielo 6 Real-time PCR system (Azure Biosystems) was used for all experiments. Total RNA was isolated using the RNeasy mini kit (Qiagen) and was digested with RNase-free DNase (Qiagen). The RNA was then reverse transcribed to cDNA using the high-capacity cDNA reverse transcription kit (Thermo Fisher) following the manufacturer's protocol.

The Power SYBR Green PCR Master Mix (Thermo Fisher) was used for real-time quantitative polymerase chain reaction (qPCR) to verify differential expression of LINE1 using primers designed on the consensus sequence of the L1MdA and L1Tf families: 5'UTR (forward: CTGCCTTGCAAGAAGAGAGC; reverse: AGTGCTGCGTTCTGATGATG), ORF1 (forward: ATCTGTCTCCCAGGTCTGCT; reverse: TCCTCCGTTTACCTTTCGCC), ORF2 (forward: GCTTCGGTGAAGTAGCTGGA; reverse: TTCGTTAGAGTCACGCCGAG), and 3'UTR (forward: AGCCAAATGGATGGACCTGG; reverse: AAGGAGGGGCATAGTGTCCA) elements[2]. GAPDH (forward: CGGCCGCATCTTCTTGTG, reverse: GTGACCAGGCGCCCAATA) was used as the normalization controls.

The TaqMan® Gene Expression Assay of JAK2 (Hs01078117_m1, ThermoFisher) was used for real-time qPCR to verify differential expression of human JAK2. Values obtained were normalized to the endogenous 18S (Hs03003631_g1, ThermoFisher) gene expression.

Relative fold changes compared to control samples was calculated by the $2^{\Delta\Delta CT}$ method. All assays were performed in triplicate.

***Human iPS cell culture and differentiation into megakaryocytes***

The BC1 iPS cell line (JAK2 wild-type, healthy donor-derived) and PVB1.4 iPS cell line (JAK2V617F homozygous, PV patient-derived) were maintained under feeder-free conditions on vitronectin-coated plates (0.5 µg/cm², Gibco, Cat. A14700) using Essential 8™ Medium (Gibco, Cat. A1517001). Cells were passaged as small clumps using 0.5 mM EDTA and maintained in an undifferentiated state. To enhance post-thaw viability, RevitaCell™ Supplement (Gibco, Cat. A2644501) was added for the first 24 hours, after which the medium was changed every 1–2 days without RevitaCell.

Hematopoietic differentiation: iPS cells were differentiated into hematopoietic cells using a modified spin-embryoid body (EB) method under feeder- and serum-free conditions[3-5]. On **Day 0**: iPS cells were dissociated into single cells using Accutase (Sigma, Cat. A6964) and seeded into 96-well untreated round-bottom plates at 3000 cells/50uL per well in StemSpan™ SFEM medium (StemCell Technologies, Cat. 09650) supplemented with RevitaCell™ (1x), 10ng/mL recombinant human FGF-2 (Peprotech, Cat. 100-18B), and 10ng/mL recombinant human BMP4 (R&D Systems, Cat. 314-BP). Plates were centrifuged at 400g for 5 minutes (no break) at room temperature to form EBs. On **Days 2, 5, 8, and 11**: 50uL of fresh StemSpan™ SFEM medium was added to each well, containing the final concentrations of 10ng/mL rhFGF-2, 10ng/mL rhBMP4, 50ng/mL recombinant human SCF (R&D Systems, Cat. 255-SC), and 10ng/mL recombinant human VEGF-A (Peprotech, Cat. 100-20) (Note: on Day 8, 100uL of media was removed from each well before adding fresh medium). On **Day 11**, 50uL of fresh StemSpan™ SFEM medium containing the final concentrations of 10ng/mL rhFGF-2, 10ng/mL rhBMP4, 50ng/mL rhSCF, 10ng/mL rhVEGF-A, and 20ng/mL recombinant human TPO (StemCell Technologies, Cat. 78210). **Day 14-18**: suspended hematopoietic cells were harvested and filtered through a 100um cell strainer (Miltenyi Biotec, Cat. 130-110-917) to remove remaining EBs.

Megakaryocytic differentiation: The harvested hematopoietic cells were cultured in StemSpan™ SFEM medium supplemented with 50ng/mL rhTPO, 10 ng/mL recombinant human IL-11 (Peprotech. Cat. 200-11), and 20 ng/mL rhSCF. Cells were seeded at $5 \times 10^5$/mL in flat-bottom, non-treated plates for 5 days, with medium replenishment on Day 3. MKs were enriched by $CD41^+$ positive selection using the EasySep Positive Selection Kit (StemCell Technologies, Cat. 17662), following the manufacturer's protocol.

***Nested allele-specific PCR for JAK2V617F-positive iPS cells***

Genomic DNA was isolated using the DNeasy Blood and Tissue Kits (Qiagen, Cat. 69504). For amplification of the *JAK2* gDNA, the PCR primers 5′-GATCTCCATATTCCAGGCTTACACA-3′ and 5′-TATTGTTTGGGCATTGTAACCTTCT-3′, which cover exon 14 of the human *JAK2* gene. The PCR product is further amplified using nested and allele-specific primers: 5'-CCTCAGAACGTTGATGGCA-3', 5'-

ATTGCTTTCCTTTTTCACAAGA-3’, 5’-GTTTTACTTACTCTCGTCTCCACAAAA-3’ (JAK2V617F mutation-specific), and 5’-AGCATTTGGTTTTAAATTATGGAGTATATG-3’ (JAK2 wild type-specific). The final PCR products were analyzed on 2.0% agarose gels. A 279-bp product indicated allele-specific JAK2V617F-positive, whereas a 229-bp product denoted allele-specific wild-type product.

***Immunohistochemistry***

Formalin-fixed, paraffin-embedded (FFPE) bone marrow sections were stained for LINE-1 ORF1p and CD41 using an adapted immunohistochemistry (IHC) protocol[6]. Slides were baked at 60℃ for 20-30min to melt paraffin, followed by sequential wash with xylene, 100%, 95%, 70%, and 50% ethanol, and running tap water for rehydration. Heat-induced antigen retrieval was performed by immersing slides in Reveal Decloaker (Biocare Medical, RV1000M, pH 6) and heating in a steamer at 98°C for 25 minutes. After cooling to room temperature, slides were washed in TBST for 5 minutes and blocked with 1% bovine serum albumin (BSA) in TBST for 30 minutes. Primary antibodies were incubated overnight at 4°C in a humidity chamber: mouse anti-human LINE-1 ORF1p (clone 4H1, Millipore Sigma, MABC1152) at 1:800 dilution (0.625 μg/mL), or rabbit anti-human CD41 (clone EPR4330, Abcam, ab134131) at 1:2000 dilution (0.325 μg/mL). After three TBST washes, endogenous peroxidase activity was quenched with 3% hydrogen peroxide for 5 minutes, followed by additional TBST washes. Secondary antibody incubation was performed for 1 hour at room temperature in a humidity chamber using IgG HRP-linked anti-mouse (Cell Signaling Technology, Cat. 7076) or anti-rabbit (Cell Signaling Technology, Cat. 7074) at 1:100 dilution. Following three TBST washes, detection was performed using the DAB substrate kit (Abcam, ab64238) with DAB enhancer (Abcam, ab675), incubating for 1 minute for both LINE-1 ORF1p and CD41. Slides were washed in water to stop the reaction, counterstained with Mayer’s hematoxylin (G-Biosciences, Cat. 786-1264), and treated with a bluing solution (10g magnesium sulfate ($MgSO_4$) and 0.67g sodium bicarbonate ($NaHCO_3$) in 1 L of deionized water). Dehydration was performed through increasing ethanol concentrations, followed by xylene. Slides were coverslipped using Limonene mounting medium (Abcam, AB104141). LINE-1 ORF1p intensity was evaluated using a four-level semiquantitative scale: 0 = negative; 1 = low; 2 = intermediate; and 3 = high.

***Quantification and statistical analysis***

Statistical analyses were performed using GraphPad Prism 9 (San Diego, CA) and *R* software. All measurements were taken from distinct samples. Except for the scRNAseq experiments, all assays were performed independently at least twice. Two-way ANOVA was used to compare blood cell counts between wild-type control and $Pf4^{+}FF1^{+}$ mice, while the two-tailed Student’s *t*-test was applied to determine statistical significance between groups. For *in vivo* experiments, randomization and blinding were not used. Sample size for animal studies were chosen to ensure a balance between the minimum numbers necessary for efficient analysis while accounting for biological variability. Figures were generated using GraphPad Prism 9, R software, and BioRender.com.
